# Spatial Multiomics Lipids and Gene expression using MALDI ISH MSI

**DOI:** 10.1101/2024.06.01.596997

**Authors:** Kyle A. Vanderschoot, Jacob P. Padilla, Kelli A. Steineman, Christopher M. De Caro, Marie C. Heffern, Elizabeth K. Neumann

**Affiliations:** Chemistry University of California, Davis; One Sheilds Dr., Davis, CA, USA

**Keywords:** in situ hybridization, single cell analysis, spatial multiomics, neuroscience, gene expression

## Abstract

Current spatial gene expression methods use DNA microarrays, Next Generation Sequencing (NGS), and fluorescence microscopy to depict the pathological/histological architecture of tissues. While each of these techniques has its own advantages, they are often costly, time intensive, and limit sampling area. A newly developed mass spectrometry-based platform, MADLI ISH MSI, combines *in-situ* hybridization (ISH) with matrix-assisted laser desorption/ionization (MALDI) to indirectly detect mRNA through an azide-modified photocleavable peptide mass tag using a single RNA targeting probe sequence. To date, 20 photocleavable mRNA probes have been synthesized to provide cellular identity within full sagittal sections of fresh frozen murine brain. This information can then be combined with existing MALDI techniques to overlay metabolomic data, such as lipids, to connect the functional state of a cell with its expressed genes. Future directions include expanding upon the number of genes that can be targeted within a single experiment.

## Introduction

Historically, transcriptomic-based measurements have been the pinnacle for understanding healthy and diseased function.^1^ Largely, this is because mRNA is the functional intermediate between DNA and expressed proteins, providing insight into both ends of the central dogma of biology. Additionally, mRNA can be sequenced and amplified, enabling a range of endogenous expression levels and different splicing variants associated with a singular gene to be concurrently measured.^2,3^ Amplification is a critical aspect that allows lowly expressed mRNA to be detected regardless of instrumental limits of detection and dynamic range, enabling high-throughout, quantitative analysis. As a result of these features, transcriptomics-based measurements have been at the forefront of biological scientific discovery.^4^ While bulk-based or single-cell measurements are informative, biological systems are composed of a unique assortment and arrangement of cells that coordinate together to enable higher order functions, such as memory and cognition within the brain.^5^ As such, there has been significant interest and development in spatially resolved transcriptomic approaches, which aim to contextualize the function of an individual cell within its cellular neighborhoods and their anatomical arrangements.^6^

To date, the main method for spatially probing gene expression is derived from *in situ* hybridization (ISH). ISH-based approaches leverage complementary base pairing between a known DNA probe and an mRNA of interest. The location of the probe and mRNA pair is generally visualized using fluorescence (FISH) and has been standard practice.^7^ Recent improvements to fluorescent microscopes has enabled subcellular, temporal resolution imaging of both live and dead cells. While incredibly powerful, fluorescence microscopy is limited to visualizing a few (usually <7) fluorescent channels at a time, due to spectral crowding and spectral bleeding/overlap.^8^ Technologies, such as MERFISH^9^, RNAScope^10^, and others circumvent this issue, by cycling fluorescent barcodes on and off of mRNA probe constructs, expanding the diversity of genes detected. Similar methodologies enable other spatial transcriptomics-based technologies, such as the Xenium platform^11^, the Geomx digital spatial profiler^12,13^, etc. To date, these systems have been widely used and demonstrate the importance of spatial biology. While important, the protocols are expensive, time consuming, and often limited in either spatial resolutions (∼50 μm) or imaging area. As a result, we have developed a method of performing *in-situ* hybridization with a mRNA-correlated mass tags, enabling the use of modern mass spectrometry imaging approaches for exploring gene expression that we have termed MALDI ISH MSI.

**Scheme 1:**
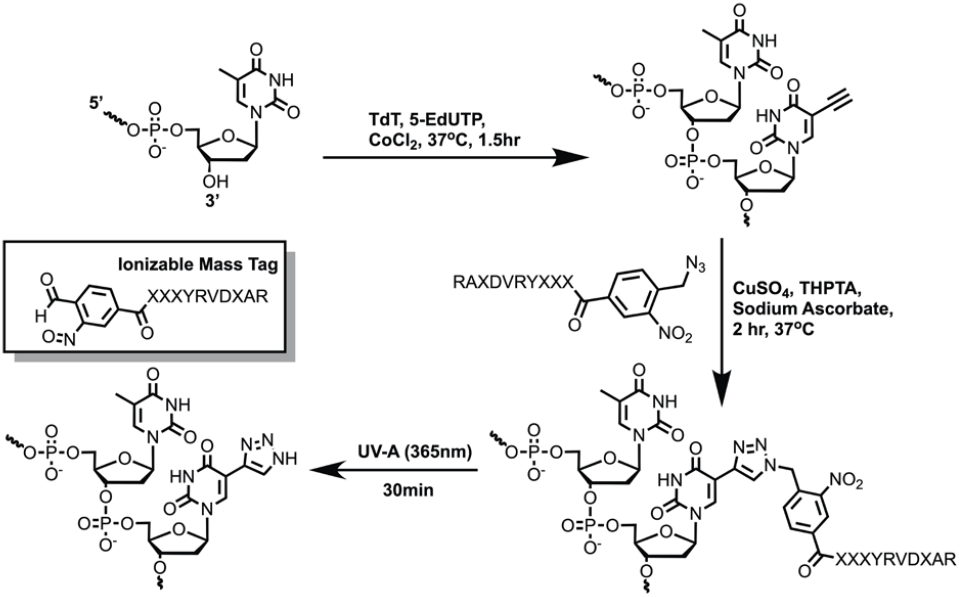
MALDI ISH probe synthesis scheme. TdT tailing with 5-EdUTP, CuAAC addition of previously synthesized azide PC mass tags and photolysis reaction products using UV-A light to yield ionizable mass tag.

Matrix-assisted laser desorption/ionization mass spectrometry imaging (MALDI MSI) is capable of measuring hundreds to thousands of molecules within a tissue without disturbing their spatial context.^14^ In brief, fresh frozen tissue is cryosectioned into 10-20 μm thick sections, thaw mounted onto a conductive surface, and coated in a UV absorbing chemical matrix to enhance and facilitate desorption and ionization of endogenous metabolites. By rastering the tissue under a UV laser, we can acquire spectra at each raster position for subsequent construction of ion images. These ion images contain information on the abundance and distribution of each detected molecule. In our case, detected mass tags are correlated to mRNA that are definitive of a cell type state or histological/pathological features. Because the main changes occur prior to MALDI MSI, this method benefits from high dynamic range, high sensitivity (attomole), speed (30 pixels/sec), mass accuracy (<5 ppm), multiplexed capacity (concurrent detection of thousands of features) and spatial resolution (<10 μm, commercially).^15^ We have also learned from other leaders in the field on various conjugation strategies for indirect detection of single mRNA and multiplexed antibodies.^16,17^ By combining these works, we have created a method capable of probe hundreds of mRNA within large sections of tissue in a cost efficient and high-throughput manner.

## Results and Discussion

Here, we describe a method of performing *in-situ* hybridization using MALDI MSI, enabling high resolution, multiplexed, multi-omic assessment of gene expression and metabolomic signatures within a single, fresh frozen tissue section (Figure 1). In brief, our method for mass labeling an mRNA probe is constructed in a flexible format (tags and probes can be mixed and matched) for affordable and scalable synthesis and application. We synthesized an *o-*nitrobenzylic azide photocleavable linker (Supplemental Figure 1) that is compatible with standard Fmoc protected peptide synthesis. Because this employs common click chemistry (specifically, CuAAC), it has high labeling efficiency.^18^ After the conjugation reaction, the resulting triazole is removed, which is less common photocage leaving group compared to *o-*nitrobenzylic alcohols or amines^19, 20^, however, enables indiscriminate chemistry. Because of the novelty, we verified photocleavage efficiency and measured side product production, through a UV cleavage test via MALDI TOF MS. The resulting photolysis products followed the expected cleavage mechanisms, yielding an aldehyde and nitroso group (Supplemental Figure 2). While the molecular ion associated with the tag was detected, there was an additional ion associated with the loss of oxygen. This has been previously observed and is noted in our sampled spectra.^21^ Because of the relatively high mass resolution of modern MALDI MSI instrumentation, this does not signficantly reduce the number of detectable labels or produce any bleedthrough as long as this loss is considered in panel design.

**Figure 1:**
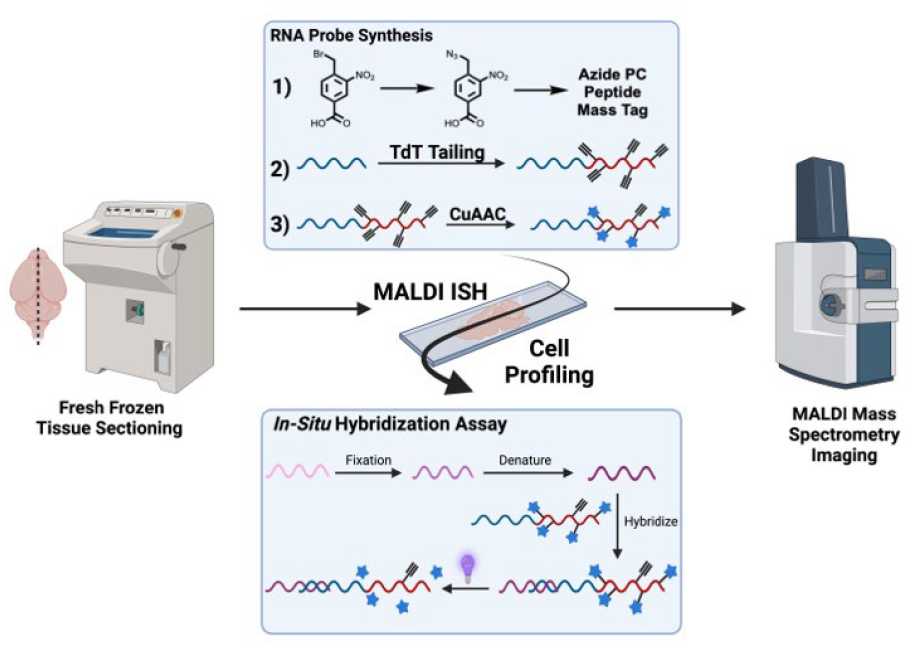
General MALDI ISH MSI workflow. Synthesized photocleavable peptide mass tags are conjugated to alkyne-tailed RNA binding probes and applied to fresh frozen tissue in a mass spectrometry-compatible *in-situ* hybridization assay. After photolytically cleaving the peptide mass tags from their RNA binding sequences, the tissue is coated in a MALDI matrix and put through a MALDI MSI experiment on a Bruker timsTOF flex.

**Figure 2:**
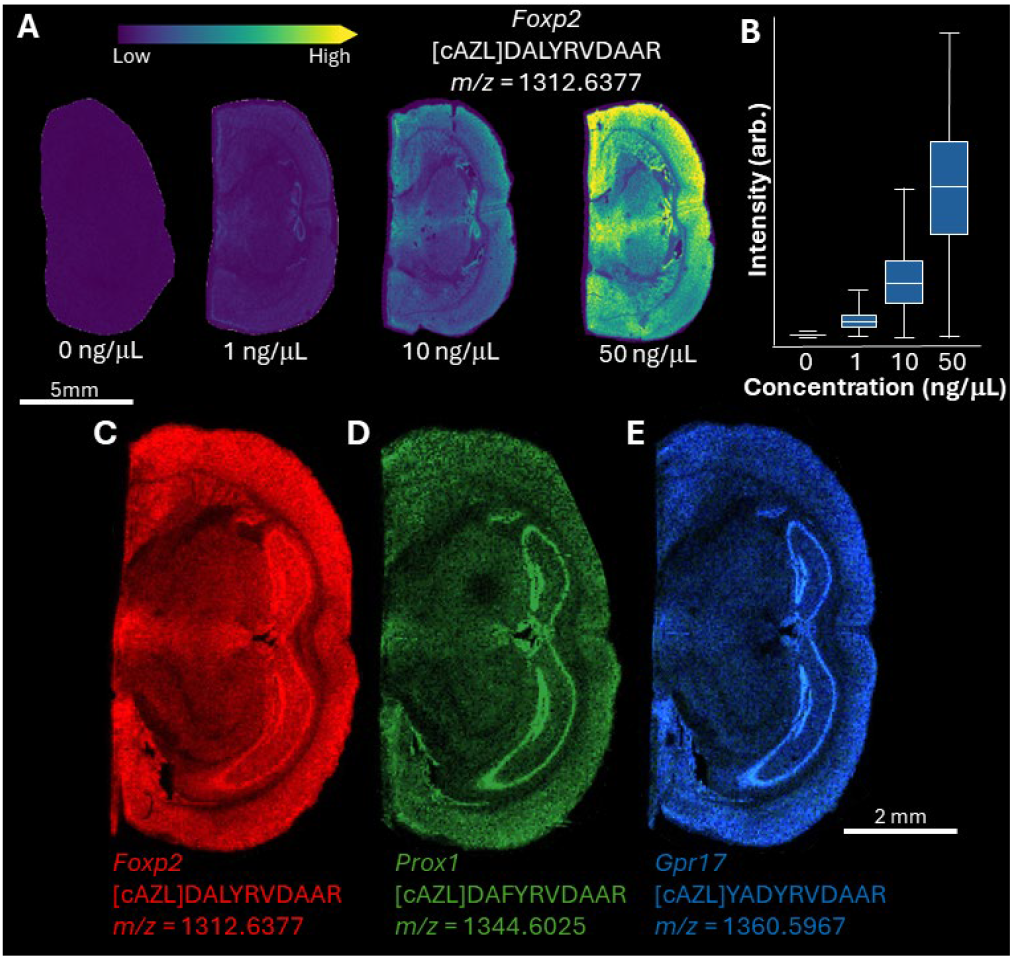
50 ng was determined to be the amount of probe necessary for detectable MS signal. A) A probe targeting mRNA associated with Foxp2 was titrated at 0 ng (blank), 1 ng/ μL, 10 ng/ μL, and 50 ng/ μL. The mass tag had a peptide sequence of DALYRVDAAR and was detected with an *m/z* value of 1312.6377. Signal was detectable at every tested concentration, although not at sufficient signal-to-noise, except at 50 ng/ μL. B) The box plot demonstrates that the expected relationship between probe amount and detected signal was achieved. C) 50 ng/ μL was sufficient for other tested probes within a different mouse brain, demonstrating that this concentration was consistent across animals and across mRNA targets.

Beyond the photocleaveable linker, we employ a peptide mass tag for detection by MALDI MSI. The peptide mass labels are easily synthesized from combinatorial peptide synthesis (from automated microwave-assited synthesizers), ensuring that each mass tag has a unique *m/z* value, can be generated in bulk, and can feature amino acids that are highly ionizable/easily detectable by MALDI MSI. By having a high ionization efficiency, we can detect the mass tags in low abundances at suffient signal-to-noise ratios. This is beneficial, because it enables the detection of a single mRNA sequencing using a single probe. In comparision, smFISH assays generally use at least 25 inidividual probes labeled in the same fluorescence channel and, even the most sophisticated spatial trancritpomics techniques require at least three probe components in the same channel.^22^ By only needing a single probe, we reduce material cost, simplify the assay’s protocol, and are capable of probing for splicing variants.^23^ Additionally, the simplified protocol, minimizes tissue degredation, enabling fresh frozen tissue to be used and does not inherently require the removal of lipids or other endogenous molecules. We accomplish this by integrating a positively charged amino acid (e.g., arginine) within the designed sequence such they have a permanent positive charge on them (Supplemental Figure 2). Although we use a peptide mass tag here, anything that can be attached to the carboxylic acid oft he photocleavable linker can be used as a mass tag, including lipids, small molecules, etc. Additionally, sequential visualization of a single tag can be done by adding, fluorescently-labeled dNTP building blocks in conjunction with the CuAAC compatible dNTP. The combination oft he two species would then allow for both fluorescence and mass spectrometry measurements on a single tissue with variable resolutions.

The RNA binding sequences were designed using LGC Biosearch Technologies’ Stellaris® Probe Designer to generate a set of 20mer sequences. The proposed sequences RNA were then filtered for GC content, homopolymer regions and potential dimerization interactions. After their synthesis, the finalized probe seuqences were then modified using template-independent tailing using terminal transferase (TdT) and 5-ethynyl-2’-deoxyuridine triphosphate (5-EdUTP). Because of the structural similarity to thymidine, the TdT enzyme is highly efficient at incorporating the click-compatible modified nucleobase to the 3’ end of the oligonucleotide. The length of the added tail was optimized (Supplemental Figure 3) to reach lengths greater than 200 bases, providing an amplification domain for the probe, allowing for single molecule dection by increasing the number of available sites for mass tag attachment. This also enables detection of mRNA using a single probe. Following the copper-catalyzed azide-alkyne cycloaddition (CuAAC), the constructed probe was purified via ethanol precipitation and put into the ISH portion of the workflow(Scheme 1).

**Figure 3:**
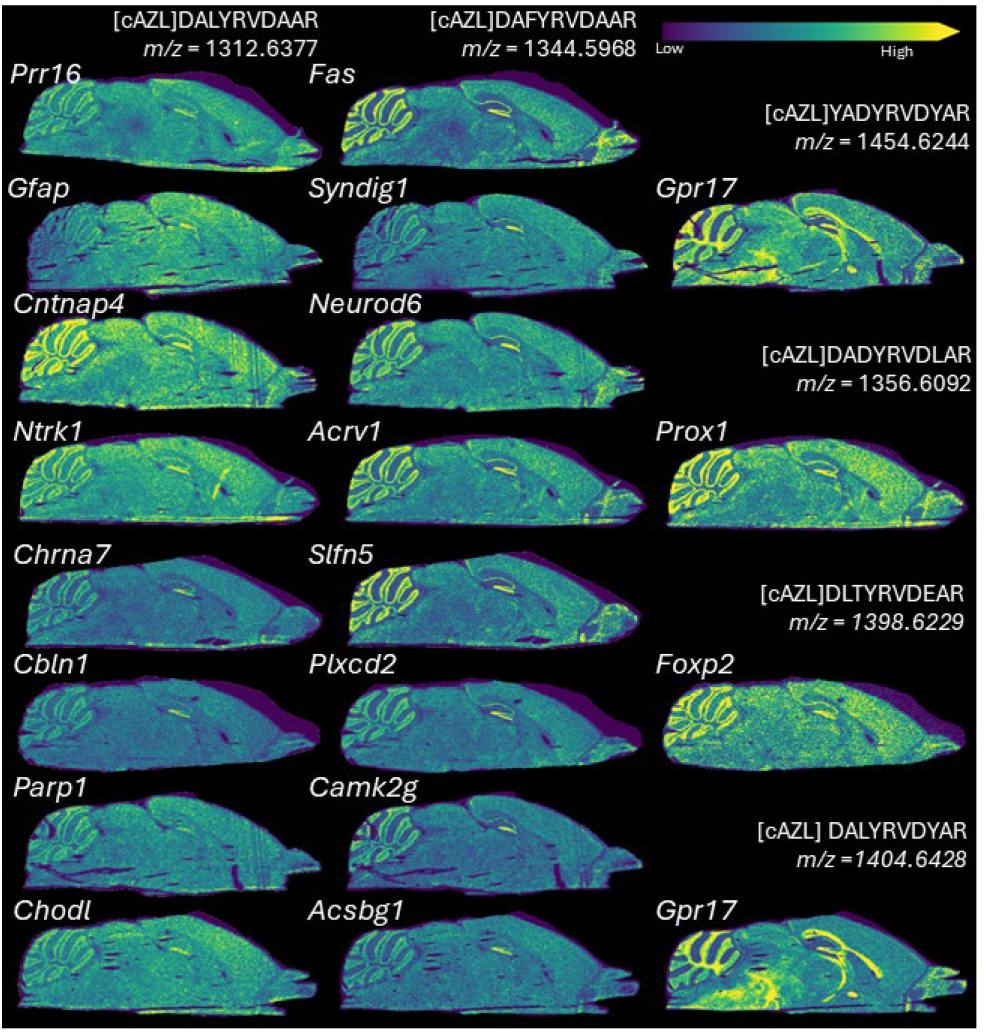
Probes were designed to target 20 mRNA relevant to neuroscience. Each of these probes were tested within sagittal sections and display high definition and signal intensity. In total 6 mass tags were tested across these 20 genes, showing probes and sequences can be exchanged. Additionally, probes were tested in sets of three, demonstrating that multiplexing is possible.

The MALDI ISH workflow follows a standard smFISH workflow^24^, with added NH_4_OAc post-hybridization washes to enable mass spectrometry analysis. In brief, the fresh frozen tissue is fixed and the RNA denatured before being incubated with the custom designed photocleavable mass tagged probes. After washing away unbound/nonspecific probes, the mass tags are photolytically cleaved with UV light (365nm), to make the mass tags available for desorption/ionization by MALDI MSI. The tissue sections are coated in a 2,5 dihydroxy acetophenone matrix and run through a MALDI imaging experiment to analyze the spatial distribution of the RNA paired with the constructed probes.

One of the key parameters associated with these measurements is the amount of probe required for adequate signal (Figure 2A), as this contributes significantly to cost. Using *Foxp2* a model probe sequence for preliminary experiments, we determinened that 50 ng/μL was sufficient for adequate spatial detection across the tissue (Figure 2A) and that this is applicable to other mRNA probe constructs: *Prox1* and *Gpr17* (Figure 2B). For a proof of concept, we show that this method is capable of visualizing twenty genes within sagittal sections of the murine brain (Figure 3). Each gene shows discrete localization throughout the sagittal section. For instance, Fas is highly expressed within granular cell layer of the cerebellum, while GFAP shows adequate expression throughout the entire section, which is expected as a generic astrocyte marker. Finally, this method enables multiomic data acquisision on one instrument. Multi-omic experiments examining native lipids and gene expression products were carried out in two separate MALDI MSI runs (Figure 4A). Lipids were first detected in negative mode, the matrix was removed and the RNA data was then gathered following the standard MALDI ISH workflow. Using this multiomics data, we demonstrate that it is possible to use the ISH profiles to label cells by identifyng genes, such as in oligodendricytes, and then extract the lipid profiles. To our knowledge, this is the first spatial lipidomics and ISH experiment performed on the same tissue section.

**Figure 4:**
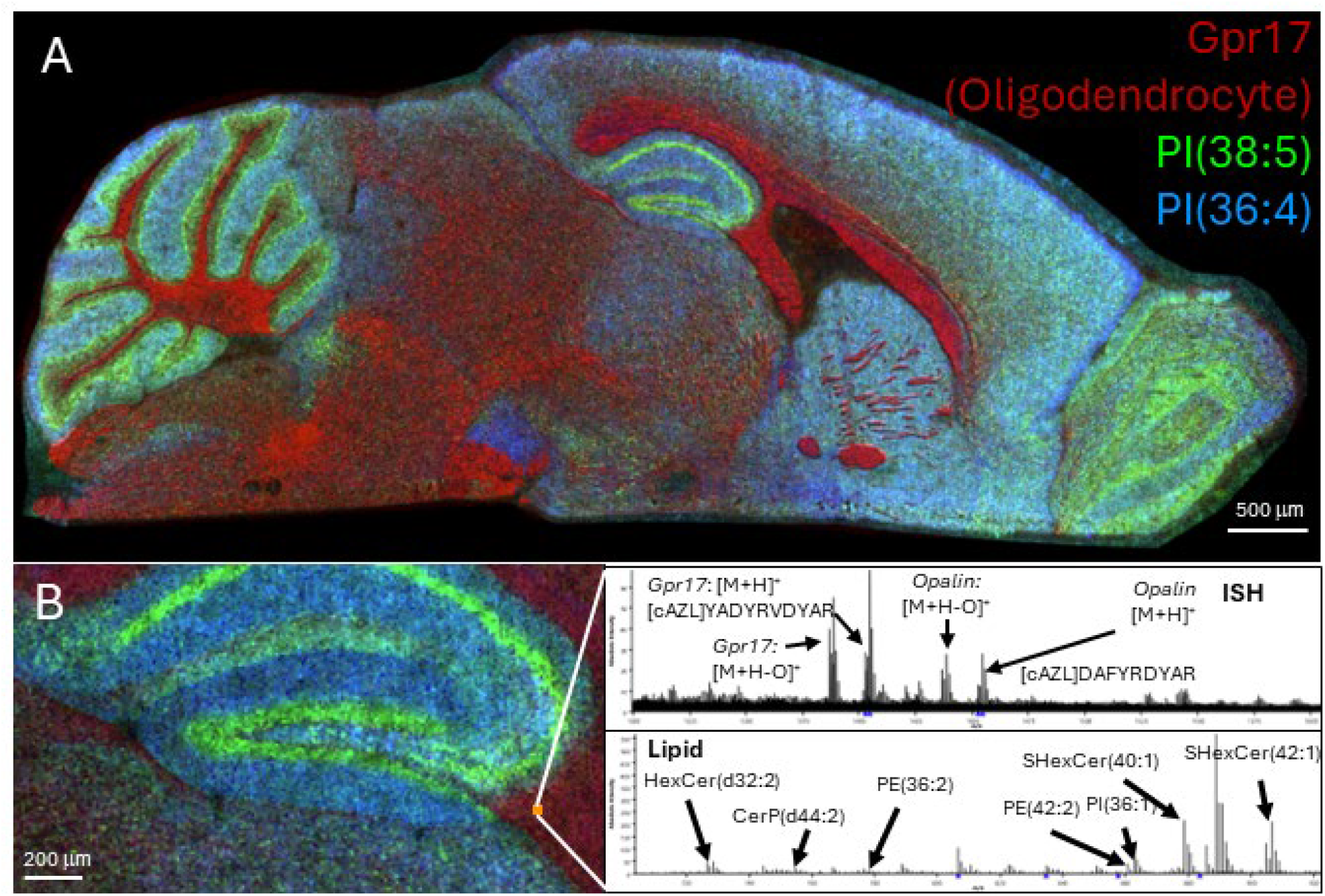
Both spatial lipidomics and gene expression were obtained from the same sagittal section, demonstrating that multiomic measurements are possible. A) MSI image of PI(38:5) (green), PI(36:4) (blue), and GPr17 (mRNA expressed in oligodendrocytes) detected within the same section. B) The lipid profile of an individual oligodendrocyte is shown, demonstrating the information that can be gained form combining spatial metabolomics with gene expression.

## Conclusion

Here, we demonstrate a method for performing MALDI ISH MSI on the murine brain for targeting 20 genes. Further, we demonstrate that lipidomic and gene expression data can be acquired on a single section to determine lipid profiles of discrete cell types. Indeed, the use of traditional click chemistries results in a flexible method that can accommodate several different mRNA targets and probe designs. The use of a mass spectrometer enables multiomic detection of lipids and mRNA features.

## Supporting Information

The authors provide additional information in the supplemental content.

## Acknowledgements

The authors gratefully acknowledge funding from the University of California, Davis. KAV acknowledges funding from the NIGMA/NIH T32 Chemical Biology Program (100007707).

